# Dear-DIA^XMBD^: deep autoencoder for data-independent acquisition proteomics

**DOI:** 10.1101/2022.08.27.505516

**Authors:** Qingzu He, Chuan-Qi Zhong, Xiang Li, Huan Guo, Yiming Li, Mingxuan Gao, Rongshan Yu, Xianming Liu, Fangfei Zhang, Tiannan Guo, Donghui Guo, Fangfu Ye, Jianwei Shuai, Jiahuai Han

**Author notes:** These authors contributed equally to this work (Qingzu He, Chuan-Qi Zhong, Xiang Li). Correspondence should be addressed to J.S. or J.H.

## Abstract

Data-independent acquisition (DIA) technology for protein identification from mass spectrometry and related algorithms is developing rapidly. The spectrum-centric analysis of DIA data without the use of spectra library from data-dependent acquisition (DDA) data represents a promising direction. In this paper, we proposed an untargeted analysis method, Dear-DIA^XMBD^, for direct analysis of DIA data. Dear-DIA^XMBD^ first integrates the deep variational autoencoder and triplet loss to learn the representations of the extracted fragment ion chromatograms, then uses the k-means clustering algorithm to aggregate fragments with similar representations into the same classes, and finally establishes the inverted index tables to determine the precursors of fragment clusters between precursors and peptides, and between fragments and peptides. We show that Dear-DIA^XMBD^ performs superiorly with the highly complicated DIA data of different species obtained by different instrument platforms. Dear-DIA^XMBD^ is publicly available at https://github.com/jianweishuai/Dear-DIA-XMBD.

## Introduction

Mass spectrometry has long been a dominant technology for peptide and protein identification and quantification^1^. The common strategy for peptide identification is performed by combining the data-dependent acquisition (DDA) approach and database search^2^. Only the top *k* peptide ions with the highest intensity are selected in an MS scan (MS1) for isolation and fragmentation in serial mode for a DDA measurement. The detected fragment ions in MS/MS spectra (MS2) are compared with the theoretical spectra generated by search engines to identify peptides. Nevertheless, the reproducibility of peptides determined by the DDA method is limited because the top *k* precursors are stochastic in repeated DDA experiments.

Aiming to overcome the limitation of the DDA mode, the data-independent acquisition (DIA) strategies have emerged, such as AIF^3^, SWATH-MS^4^, HDMS^E5^, MSX^6^, WiSIM-DIA^7^, SONAR^8^, HRM^9^, BoxCar DIA^10^, diaPASEF^11^, Scanning SWATH^12^ and PulseDIA^13^. A common DIA mode is named sequential window acquisition of all theoretical mass spectra (SWATH-MS), in which all peptide ions in a specified isolation window with a large mass-to-charge ratio (m/z) are fragmented. The mass spectrometer records all the fragment signals of the mixed peptides in an isolation window. Obviously, it is extremely difficult to directly analyze DIA data because the peptide and fragment signals are mixed in corresponding MS and MS/MS spectra.

In recent years, a number of methods have been developed to process DIA data. For instance, the library-based tools include Spectronaut^9^, OpenSWATH^14^, SWATHProphet^15^, Skyline^16^, Specter^17^, EncyclopeDIA^18^, PIQED^19^, DIA-NN^20^, and MaxDIA^21^; the library-free tools include DIA-Umpire^22^, Group-DIA^23, 24^, directDIA (a part of Spectronaut), MSPLIT-DIA^25^, PECAN^26^, DeepNovo-DIA^27^, and MaxDIA; and the library predicting tools contain DeepMass^28^, pDeep^29^, Prosit^30^, and DeepDIA^31^. OpenSWATH, a prevalent library-dependent workflow integrated into OpenMS^32^, was proposed to analyze the SWATH-MS data. OpenSWATH scores the peptides in SWATH-MS data based on the spectral library typically built on DDA MS. In order to overcome the limitation of DDA library generation, DIA-Umpire calculates the correlation coefficient between precursors and fragments to generate the pseudo-DDA spectra. Group-DIA analyzes the multiple DIA data files simultaneously to determine the precursor-fragment pairs. Both DIA-Umpire and Group-DIA are based on the spectrum-centric strategy. PECAN is a peptide-centric analysis tool that requires a peptide sequence-based library to directly detect peptides from DIA data. MSPLIT-DIA employs the peptide query method to analyze each DIA MS/MS spectrum. However, the conventional statistical algorithms used by these DIA methods make them insufficient for pattern recognition and classification of fragment XICs.

In the past two years, several deep learning-based methods have been developed to analyze DIA data^33, 34, 35^. DeepNovo-DIA combines the de novo peptide-sequencing method and deep learning to directly identify the amino acid sequences from DIA spectra. DIA-NN begins with a peptide-centric strategy based on *in silico* spectra libraries and then uses a deep neural network to calculate the q-value of peptides. DeepDIA predicts MS/MS spectrum and the normalized retention time (iRT) of peptides in a protein database with a deep learning model and then generates *in silico* spectral libraries to analyze DIA data. Nevertheless, none of the above-mentioned deep-learning-based methods directly analyze DIA data to produce tandem spectra for database-searching and then generate the internal libraries (DIA-derived) for quantification. In addition, all of these methods apply supervised learning methods, which limits their generalization ability.

In this paper, we developed Dear-DIA^XMBD^, a spectrum-centric method that combines the deep variational autoencoder^36^ (VAE) with other machine learning algorithms to detect the correspondence between precursors and fragments in DIA data without the help of DDA experiments. Dear-DIA^XMBD^ produces the pseudo-tandem MS spectra to a search database and generates the internal libraries. Our approach can be easily integrated into the existing workflow because the output file of Dear-DIA^XMBD^ is in MGF format that can be processed by common search engines, including Comet^37^, X!Tandem^38^, and MSFragger^39^. Furthermore, benefiting from the fact that the autoencoder is an unsupervised deep learning model, Dear-DIA^XMBD^ shows excellent performance on the DIA data of different species obtained by different instrument platforms. Due to its powerful generalization ability, we suggest that Dear-DIA^XMBD^ is a valuable open-source software for DIA proteomics.

## Results

### Dear-DIA^XMBD^ workflow

DIA data is usually visualized as three-dimensional data containing m/z, retention time, and intensity. In order to correctly link the precursors and the ions produced, Dear-DIA^XMBD^ first splits the MS1 retention time with fixed-width sliders in each isolation window. Each precursor slider is treated as a minimum processing unit containing a series of MS1 spectra and corresponding MS2 spectra (**Fig. 1a**).

**Figure 1.**
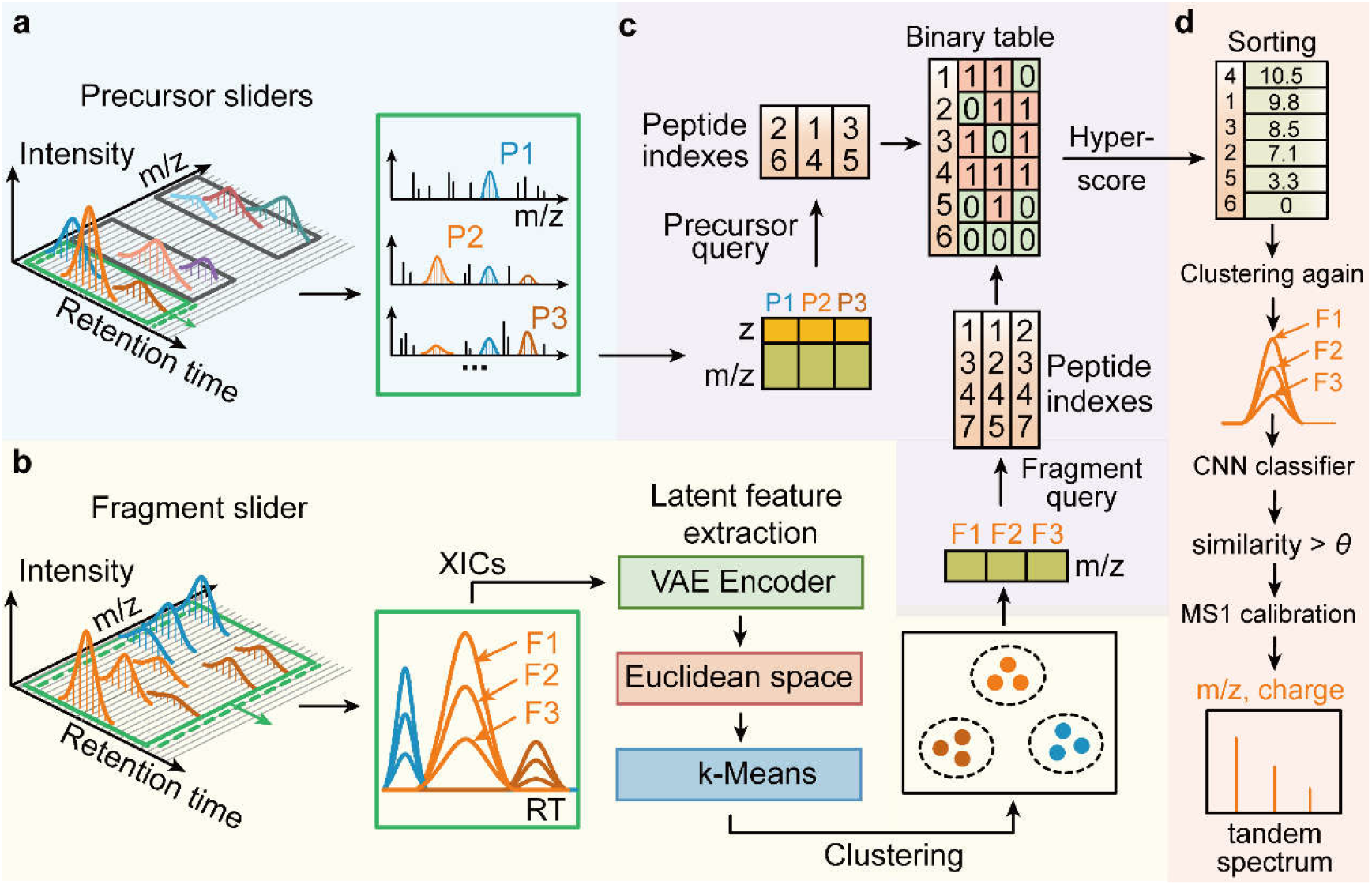
The workflow of Dear-DIA^XMBD^. **(a)** A precursor slider advanced using set strides along the retention time dimension. The candidate precursors (described in the Methods section) are detected by several SNR-dependent algorithms. **(b)** The candidate fragment XICs (described in the Methods section) are embedded into the Euclidean space after being fed to the VAE encoder neural network and then assigned to k-classes using a k-Means clustering algorithm. **(c)** Each fragment cluster is combined with the corresponding precursor based on the protein database and hyper-score. **(d)** After the high-scoring precursor fragment pairs are removed, the remaining ions are clustered again by using k-Means. These precursor-fragment pairs are judged using a convolutional neural network (CNN) to calculate the similarity among fragments matching the *in-silico* spectrum. The precursor-fragment groups with high similarity are stored as pseudo-tandem spectra for identification.

Next, we removed the background ions with a low signal-noise ratio (SNR) in the slider by using the peak-finding and deisotoping algorithms to determine the candidate precursors and fragments (described in the **Methods** section). Since the point-to-point similarity calculation between XICs of candidate fragments is affected by noise and peak misalignment, we employed the VAE encoder (see **Supplementary Fig. 1** for details) to extract the latent features of fragment XICs and then mapped these features to the Euclidean space. Then the k-Means clustering algorithm using a Euclidean metric is applied to assign the candidate fragments to k classes in the feature space (**Fig. 1b**).

Ideally, the fragments in the same cluster should be from the same precursor. In our model, we provide a peptide indexing algorithm named PIndex, which is designed for closed-search to return the unique indexes of *in silico* digested peptides obtained from the FASTA database to determine the precursor of each fragment cluster. A binary table presents the intersection of two peptide index sets, that is, the precursor index set obtained from the theoretical identity of the precursor query and the fragment index set obtained from the theoretical identity of the fragment query (**Fig. 1c** and the **Methods** section).

We then calculated the hyper-score and sorted the scores for all the *in silico* digested peptides based on the binary table. Afterward, we removed the precursor-fragment pairs with high scores in the clustering results and then performed k-Means clustering again on the remaining ions (**Fig. 1d** and the **Methods** section). A convolutional neural network (CNN) (**Supplementary Fig. 2** and the **Methods** section) was applied to calculate the similarity among the sets of fragments matching the highest score *in silico* digested peptide. If the similarity exceeds a certain threshold (*0*), the fragments in each cluster will be grouped with the corresponding precursor. We used the high score precursor-fragment groups as internal calibrants to recalibrate all the precursors m/z^40^. Finally, the calibrated precursor-fragment pairs were stored as the tandem spectrum (**Fig. 1d** and the **Methods** section).

### Applications of deep VAE and inverted index

Due to the high-order complexity of DIA data, the direct classification of the mixed and unlabeled fragment XICs is extremely difficult. Therefore, we designed a VAE model comprised of encoder and decoder networks for classification (**Fig. 2a**). The principle of this model is that the triplet fragment XICs are entered into a multi-branched encoder network to extract the latent features of input data for cluster analysis in Euclidean space (**Fig. 2a**). During the training process, the latent features are reconstructed by a decoder network to make them as close as possible to the input of the encoder (**Fig. 2a** and **Supplementary Note 1**). We employed a loss function of classical VAE with the triplet loss function of FaceNet^41^ to improve the ability of the model to distinguish fragments from different precursors. Using a combination of the triplet loss and VAE, we can generate similar representations for fragments of the same precursor and produce the dissimilar features for fragments of the different precursors (see the **Methods** section, **Supplementary Fig. 1** and **Supplementary Note 1** for details).

**Figure 2.**
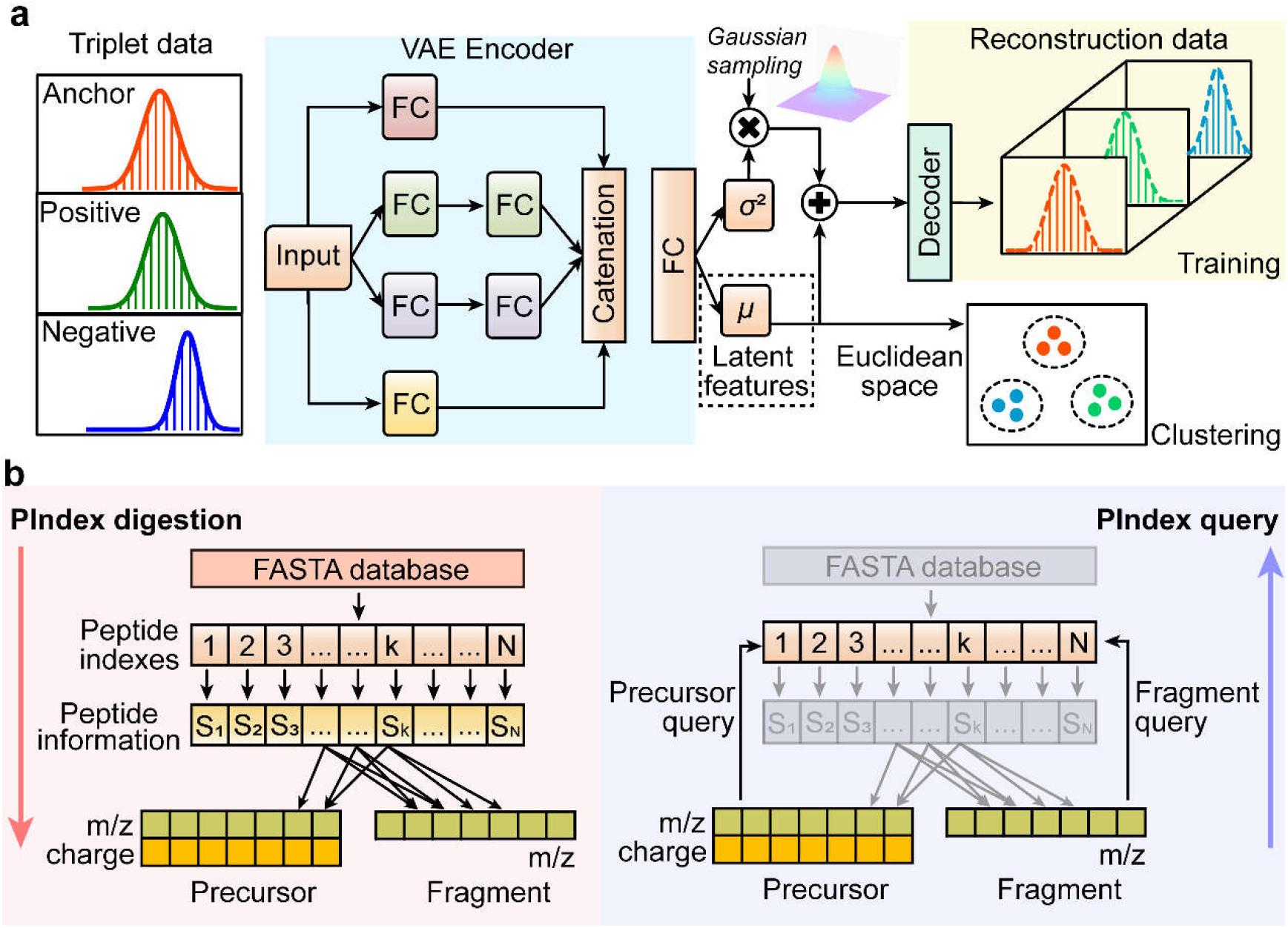
The schematic diagrams of the Deep Leaning model and PIndex querying algorithm. **(a)** The structure of the deep variational autoencoder (VAE). The input data contains three components: the anchor-fragment XIC (red), the positive (green), and the negative (blue) fragment XICs (see the Methods section for details). The anchor and positive fragments come from the same peptide; meanwhile, the anchor and negative fragments belong to different peptides. The triplet data is fed to the four-branch encoder network, which is consisted of 1-2-2-1 fully connected (FC) layers. The output vectors of the four-branch networks are catenated by the appending operation at the end. The encoder network outputs two vectors of equal size, one for the variance (σ^2^) and the other for the mean value (μ). The mean vector represents the latent features of the input data. Since anchor and positive are from the same peptide but anchor and negative are from different peptides, the anchor fragment is closer to the positive fragment than to the negative fragment after training triplet loss (see the Methods section and Supplementary Note 1 for details). **(b)** The peptide indexing algorithm (PIndex). The left part shows the protein database digested into a variety of sets Sk, where k indicates the unique index of peptides. The right part describes the processes of precursor query and fragment query (described in the Methods section). The peptide indexes can be queried by m/z-charge pairs of precursors and m/z of fragments, respectively.

Next, we addressed how to precisely match the precursor and fragment clusters. As the closed-search is still the main strategy of a database search engine such as Comet and X!Tandem, an indexing algorithm named PIndex is used to connect the clustering results with the candidate precursors (described in the **Methods** section). PIndex contains PIndex digestion and PIndex query algorithms. PIndex digestion begins with an *in silico* digestion of the protein database containing the series of sets of peptide information Sk with each Sk corresponding to a unique peptide index k (**Fig. 2b**). A peptide information set contains the charge of precursor, the m/z of precursor, and the m/z list of fragments. To determine the precursor of each fragment cluster, PIndex constructed precursor query^42^ and fragment query for *in silico* precursors and fragments in all peptide information sets, respectively (see the **Methods** section for details). Precursor and fragment queries apply the m/z-charge pairs and the m/z ratios as the key for querying peptide indexes, respectively (**Fig. 2b**). With the peptide indexes as relay stations, the precursor of each fragment cluster can be quickly inferred.

### A comparison of Dear-DIA^XMBD^ with other DIA analyzing software

To evaluate the performance of Dear-DIA^XMBD^, we make a comparison of Dear-DIA^XMBD^ with the available DIA analysis approaches of DIA-Umpire and Spectronaut 14. First, we trained the autoencoder of Dear-DIA^XMBD^ on an *E. coli* SWATH dataset with 100 variable MS1 windows, which are acquired by TripleTOF 5600 mass spectrometer and TripleTOF 6600 mass spectrometer. The dataset obtained from TripleTOF 5600 contains seven runs with the MS recording time varying from 30 minutes to 240 minutes. The datasets from TripleTOF 6600 consist of six runs with the MS recording time varying from 15 minutes to 10 hours^43^. We manually selected 97,980 *E. coli* peptide precursor ions quantified by OpenSWATH. Each precursor ion contains the top 6 fragment ion XICs. Then, we randomly picked two fragment XICs of the same precursor ion as an anchor and positive XICs, and randomly selected a fragment XIC from other precursor ions as negative XIC to generate a total of 2179,590 groups of triplet data as the training dataset (**Supplementary Note 1**). Different from the common supervised deep learning models, we applied the autoencoder to extract the characteristics of XICs, which allows us to use only the number of quantified proteins and peptides as indicators to optimize the model.

We benchmarked the performance of Dear-DIA^XMBD^ by using the highly complicated sample datasets, which consist of SWATH-MS Gold Standard (SGS) human dataset^14^, L929 mouse dataset, and HYE124 dataset^44^ with 64 variable windows (AB Sciex TripleTOF 6600). We employed Dear-DIA^XMBD^ to generate pseudo-tandem spectra and then combined Comet and X!Tandem (native and k-score version) to search the protein FASTA database for peptides and protein identification. All identified peptides and proteins were filtered with a 1% false discovery rate (FDR) of peptide-level score determined by MAYU^45^ to establish the spectrum libraries, and then OpenSWATH-PyProphet-TRIC workflow was applied to quantify peptides and proteins in libraries from Dear-DIA^XMBD^ (**Supplementary Fig. 3**). We applied DIA-Umpire to generate pseudo-tandem spectra and then used the same software tools to process the tandem-spectra file. In addition, we also used Spectronaut 14 (directDIA 2.0) to analyze the benchmark datasets and set the 1% Q-value for filtering the peptides and proteins. In the reports of Dear-DIA^XMBD^, Spectronaut 14, and DIA-Umpire, we removed the proteins with only one peptide found and compared the proteins with two or more peptides found.

The SGS human dataset was generated by Röst *et al*. by using the separately diluted 422 stable isotope-labeled standards (SIS) peptides in HeLa cell lysate in 10 dilution steps (from 1× dilution to 512× dilution), and then acquired as DIA data in triplicate with SWATH-MS. According to the quantified results of SIS peptides, Dear-DIA^XMBD^ can find more synthesized peptides than Spectronaut 14 and DIA-Umpire in all dilution steps, indicating that the sensitivity of Dear-DIA^XMBD^ is higher than Spectronaut 14 and DIA-Umpire (**Fig. 3a**). Dear-DIA^XMBD^ covers 97% and 98% (average coverage) of SIS peptides reported by Spectronaut 14 and DIA-Umpire, respectively. Notably, the number of SIS peptides uniquely discovered by Dear-DIA^XMBD^ far exceeds those found by Spectronaut 14 and DIA-Umpire, demonstrating that Dear-DIA^XMBD^ shows a higher confidence interval (**Fig. 3a**). In addition, Dear-DIA^XMBD^ finds more human peptides and proteins than Spectronaut 14 and DIA-Umpire when analyzing 10 dilution steps combined data. According to the quantified results, Dear-DIA^XMBD^ discovered 23,141 peptides and 2,616 proteins, while DIA-Umpire reported 11,080 peptides and 1,254 proteins, and Spectronaut 14 found 17,579 peptides and 1,928 proteins. (**Fig. 3b** and **Supplementary Fig. 4**).

**Figure 3.**
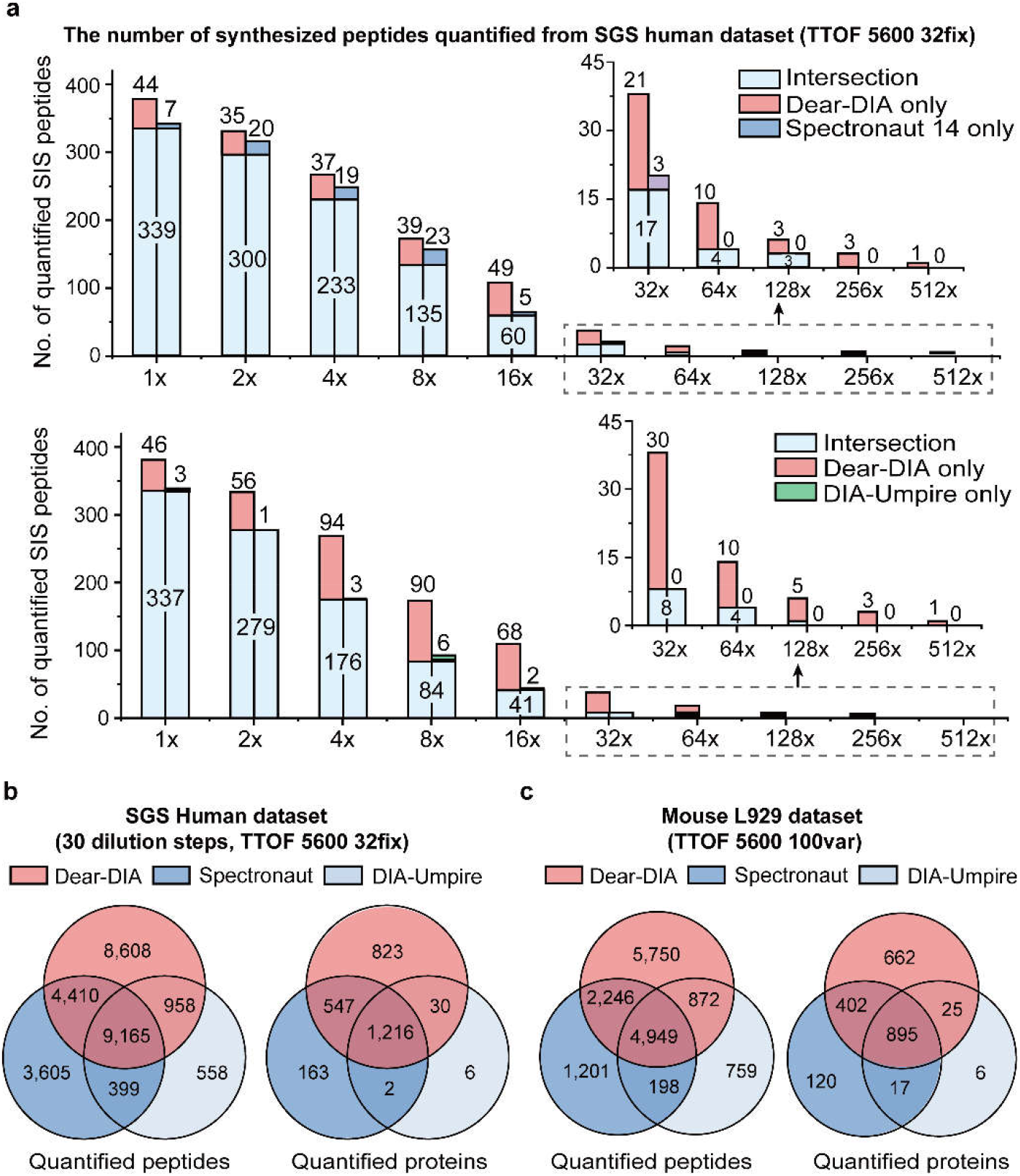
Analysis results of SGS human dataset and mouse L929 dataset. **(a)** The number of synthesized peptides from the SGS human dataset. The horizontal axis shows the dilution steps from 1× dilution to 512× dilution. The light blue parts in the upper and lower histograms represent the intersection of Dear-DIA^XMBD^ and Spectronaut 14, and the intersection of Dear-DIA^XMBD^ and DIA-Umpire, respectively. The red, dark blue and green parts show the stable isotope-labeled standard (SIS) peptides uniquely found by Dear-DIA^XMBD^, Spectronaut 14, and DIA-Umpire, respectively. **(b)** Analysis results of all dilution steps (from 1× dilution to 512× dilution), total 30 files, in SGS human dataset. The red, the dark blue and the light blue circles represent the results of Dear-DIA^XMBD^, Spectronaut 14, and DIA-Umpire, respectively. **(c)** Venn diagrams of peptides and proteins found using mouse L929 dataset. The red, the dark blue, and the light blue circles represent the results of Dear-DIA^XMBD^, Spectronaut 14, and DIA-Umpire, respectively.

The mouse dataset was derived from L929 cell lysate, which contains triplicate samples with 100 variable MS1 windows measured in SWATH mode on TripleTOF 5600 mass spectrometer (AB Sciex). In the quantification process, the total numbers of peptides found by Dear-DIA^XMBD^, Spectronaut 14, and DIA-Umpire were 13,817, 8,594, and 6,778, respectively, and the total number of proteins found by Dear-DIA^XMBD^, Spectronaut 14, and DIA-Umpire were 1,984, 1,434, and 943, respectively. Dear-DIA^XMBD^ also covers 83% of peptides and 90% of proteins reported by Spectronaut 14. Dear-DIA^XMBD^ covers 87% of peptides and 98% of proteins revealed by DIA-Umpire. The wide coverage shows a nice reproducibility among Dear-DIA^XMBD^, DIA-Umpire, and Spectronaut 14 (**Fig. 3c** and **Supplementary Fig. 4**). Dear-DIA^XMBD^ discovered more low-intensity peptides than DIA-Umpire (**Supplementary Fig. 9-10**).

Next, we compare the performances of Dear-DIA^XMBD^, DIA-Umpire, and Spectronaut 14 with the HYE124 dataset, which was specifically designed for checking DIA algorithms. The HYE124 dataset includes two hybrid proteome samples, A and B. Sample A was composed of 65% w/w human, 30% w/w yeast, and 5% w/w E. coli proteins, while sample B was composed of 65% w/w human, 15% w/w yeast, and 20% w/w E. coli proteins.

Adding two samples of HYE124 datasets together, the total quantified peptides discovered by Dear-DIA^XMBD^, Spectronaut 14, and DIA-Umpire are 63,142, 51,813, and 33,016, respectively, and the total quantified proteins are 6,717, 5,025, and 3,242, respectively, in which Dear-DIA^XMBD^ covers 93% proteins and 78% peptides found by Spectronaut 14. Dear-DIA^XMBD^ also covers 99% proteins and 86% peptides found by DIA-Umpire (**Fig. 4a**). These results show quite a good reproducibility among Dear-DIA^XMBD^, Spectronaut 14, and DIA-Umpire. In addition, the number of identified peptides discovered uniquely by Dear-DIA^XMBD^ was 7.9 times that found uniquely by DIA-Umpire (i.e., 37,681 versus 4,768) (**Fig. 4a**). Dear-DIA^XMBD^ can find a large number of proteins and peptides overlooked by DIA-Umpire in identification and quantification. The current Dear-DIA^XMBD^ only uses the *E. coli* data as training data, but it shows excellent generalization ability.

**Figure 4.**
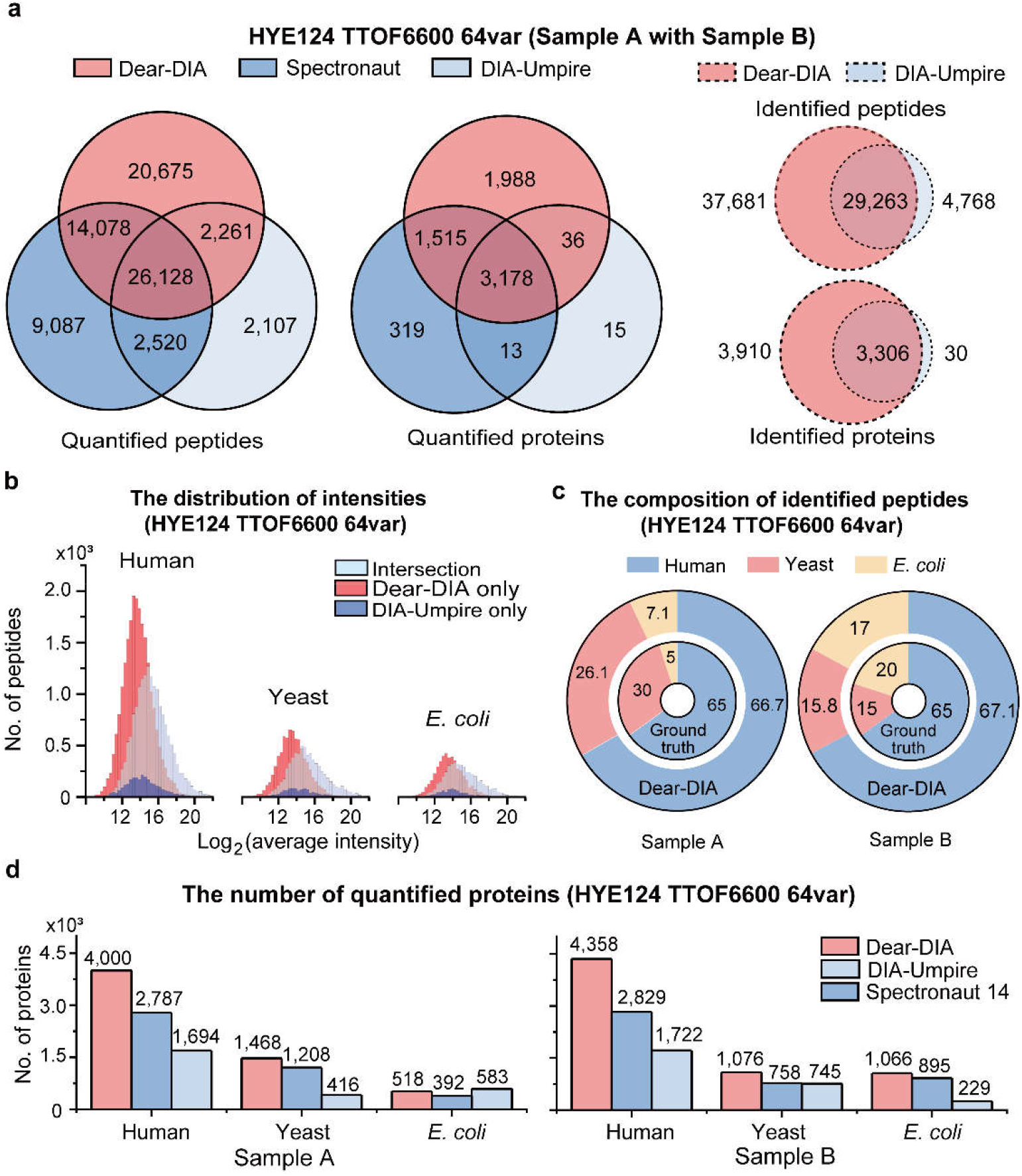
Analysis results of HYE124 dataset with 64 variable windows (TripleTOF 6600). **(a)** The comparison of numbers of identified and quantified peptides and proteins obtained by Dear-DIA^XMBD^, DIA-Umpire, and Spectronaut 14 from the HYE124 dataset with samples A and B together. The solid lines and the dashed lines show the quantified and identified results, respectively. The red, the dark blue, and the light blue circles represent the results of Dear-DIA^XMBD^, Spectronaut 14, and DIA-Umpire, respectively. **(b)** The log2-scaled distributions of the quantified peptide intensities were discovered from the HYE124 dataset with samples A and B together. The peptides shared jointly with DIA-Umpire and Dear-DIA^XMBD^ are shown in light blue; the peptides exclusively reported by Dear-DIA^XMBD^ and by DIA-Umpire are shown in red and dark blue, respectively. **(c)** The composition of proteins found by Dear-DIA^XMBD^ with sample A dataset and with sample B dataset, respectively. The blue, red, and yellow colors represent human, yeast, and *E. coli* species, respectively. The smaller doughnuts represent the ground-truth composition of proteins, which are mixed in defined proportions. The larger doughnuts show the composition of proteins discovered by Dear-DIA^XMBD^. **(d)** The numbers of the quantified proteins found by Dear-DIA^XMBD^, Spectronaut 14, and DIA-Umpire with sample A dataset and sample B dataset, respectively. The red, dark blue and light blue colors show the results of Dear-DIA^XMBD^, Spectronaut 14, and DIA-Umpire, respectively.

Furthermore, it is well known that proteins and peptides with low abundance are hardly identified by MS analysis algorithms because of the interference of background noise. However, Dear-DIA^XMBD^ performs much better than DIA-Umpire on this issue when using the same quantified software tool such as OpenSWATH since the intensity distributions of the quantified proteins and peptides given by Dear-DIA^XMBD^ are more in the low-density range (**Fig. 4b** and **Supplementary Fig. 11-12**).

We also analyzed sample A and sample B of the HYE124 dataset separately. For sample A, the percentages of identified peptides given by Dear-DIA^XMBD^ were 66.7% for humans, 26.1% for yeast, and 7.1% for *E. coli*, respectively. For sample B, Dear-DIA^XMBD^ found 67.1% of humans, 15.8% of yeast, and 17% of *E. coli* identified peptides, respectively (**Fig. 4c**). Consistently, Dear-DIA^XMBD^ found more proteins in humans, yeast, and *E. coli* than Spectronaut 14 and DIA-Umpire (**Fig. 4d**). We manually checked the XICs of human, yeast, and *E. coli* peptides identified by Dear-DIA^XMBD^ (but not DIA-Umpire and Spectronaut 14) to confirm the similarity among fragments (**Supplementary Fig. 13**).

We used LFQbench^44^ R package to benchmark the precision of quantification on the HYE124 dataset. Compared with Spectronaut 14 and DIA-Umpire, Dear-DIA^XMBD^ performs similarly in precision for both peptides and proteins of humans, yeast, and *E. coli* (**Fig. 5, Supplementary Fig. 14 and Supplementary Table 1**). We also tested the performance of Dear-DIA^XMBD^ on the HYE124 TripleTOF 5600 dataset with 64 variable windows (**Supplementary Fig. 15-18**). Dear-DIA^XMBD^ discovered more peptides and proteins than Spectronaut 14 and DIA-Umpire.

**Figure 5.**
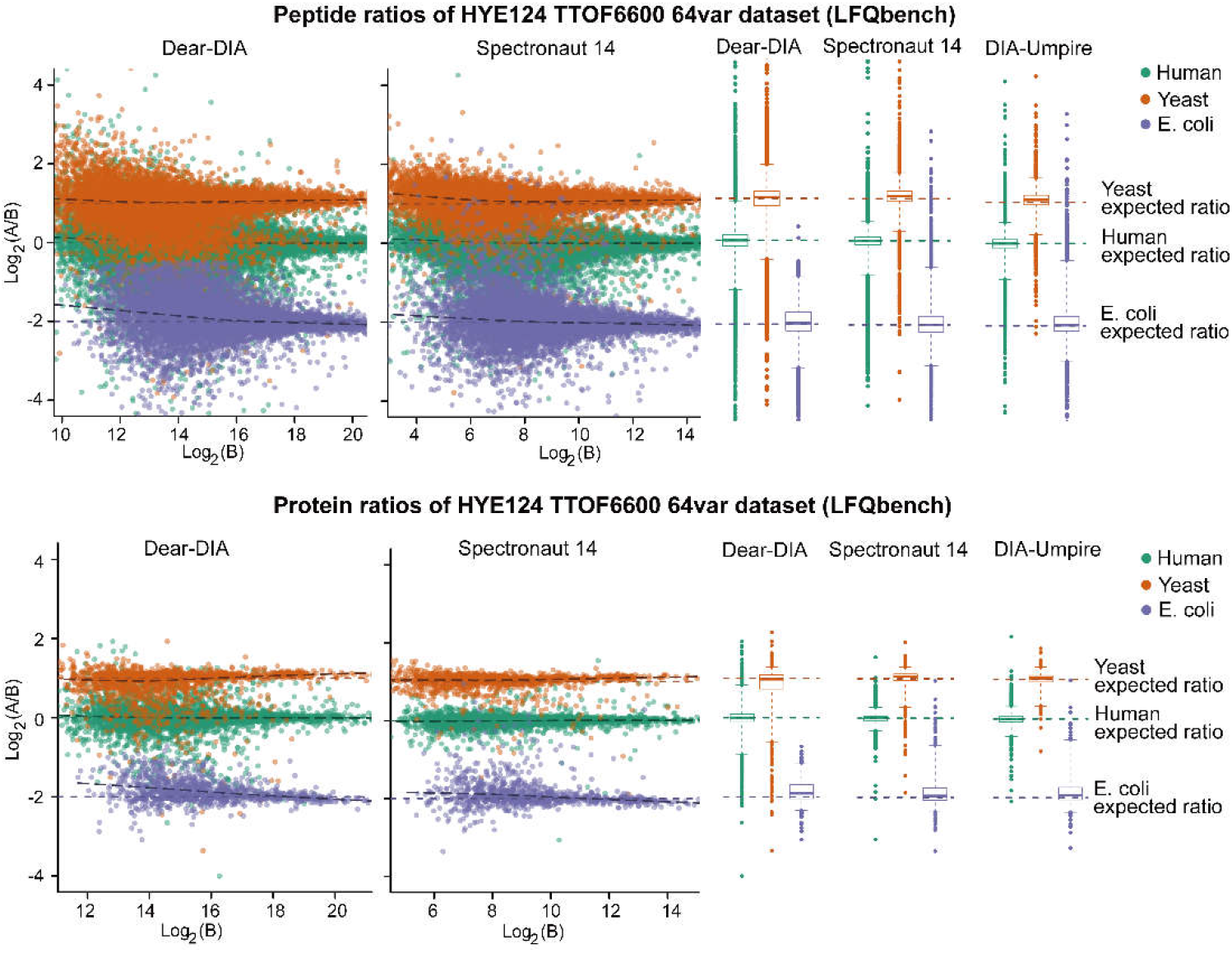
Peptide-level and protein-level LFQbench test performance of HYE124 dataset with 64 variable windows (TripleTOF 6600). The upper and lower scatter plots represent the peptide ratios and the protein ratios reported by Dear-DIA^XMBD^ and Spectronaut14, respectively. The colored dashed lines indicate the expected log_2_(A/B) ratios for human (green), yeast (orange), and *E. coli* (purple) species. The black dashed lines represent the local trend along the x-axis of the experimental log-transformed ratios of each population (human, yeast, and E. coli). The horizontal axis and vertical axis of the scatter chart represent the log-transformed ratios (log_2_(A/B)) of the quantified intensity and the log-transformed intensity of sample B (log_2_(B)), respectively. The upper and lower box plots show the quantified performance of peptides and proteins, respectively (boxes, interquartile range; whiskers, 1–99 percentile; human, yeast, and *E. coli*).

Additionally, we used Dear-DIA^XMBD^, Spectronaut 14, and DIA-Umpire to analyze the Biognosys facility (BGS) mouse DIA dataset^46^, which was acquired from Orbitrap Fusion Lumos mass spectrometer (Thermo Fisher Scientific, San Jose, CA) with a 2-hour gradient and 40 DIA scans. The results demonstrated that Dear-DIA^XMBD^ could also analyze the data from Thermo Fisher Scientific mass spectrometer (**Supplementary Fig. 19**).

We also benchmarked Dear-DIA^XMBD^ on TNFR1 (tumor necrosis factor receptor 1) complex dataset^23, 47, 48, 49^ from L929 cells treated with TNF from six different time periods. We performed a comparison of OpenSWATH quantified results and manual inspection results, showing that Dear-DIA^XMBD^ can find truly regulated proteins (**Supplementary Fig. 20**).

## Discussion

In the paper, we designed a new method with neural network architecture, namely Dear-DIA^XMBD^, to improve the feature extraction ability of fragment XICs, which consults to the structures of a fully connected VAE network. Moreover, we also implemented a high concurrency program written in C/C++ from scratch to increase the speed of the program.

We demonstrated that Dear-DIA^XMBD^ is a more efficient method than DIA-Umpire and Spectronaut 14 in directly analyzing DIA data to discover proteins and peptides. First of all, Dear-DIA^XMBD^ can reproduce most of the results obtained by DIA-Umpire and Spectronaut 14 at identification and quantification levels. Secondly, Dear-DIA^XMBD^ can identify more low-abundance proteins and peptides, indicating that it has a better performance than DIA-Umpire in processing the low SNR signals.

Furthermore, although the training dataset is from *E. coli*, Dear-DIA^XMBD^ shows an outstanding performance in analyzing datasets of different species with different instruments, indicating its general recognition ability. The pseudo-tandem spectra generated by Dear-DIA^XMBD^ can be easily fed into common search engines for library generation. Additionally, analyzing mass spectrometry data of post-translational modifications (PTMs) is an important issue and challenge. Dear-DIA^XMBD^ currently supports carbamidomethyl as a fixed modification and oxidation and n-acetylation as variable modifications. Dear-DIA^XMBD^ does not work accurately enough for other modifications such as phosphorylation (see details in **Supplementary Note 4**). Since OpenSWATH IPF^50^ is a powerful tool for processing PTMs data, we consider combining Dear-DIA^XMBD^ and OpenSWATH IPF to analyze PTMs DIA data in future work.

Collectively, Dear-DIA^XMBD^ is an advanced software for processing a variety of highly complex DIA data. We believe that Deep Learning methods may play more important roles in the analysis of the complicated protein spectrum data.

## Acknowledgments

We would like to express our deep appreciation to Dazheng Wang, Steven X. Shuai, Peter Shaw, Helen Shaw and Xiaojie Liu for the helpful suggestions while drafting the manuscript. We thank Z. Xu and Y. Yu for help with the high-performance computer.

## Author Contributions

Jianwei Shuai, Qingzu He, and Chuan-Qi Zhong conceived the project. Qingzu He developed the algorithm and implemented the software, and wrote the manuscript. Chuan-Qi Zhong acquired mass spectrometry data for training the deep neural network. Xiang Li analyzed the data and results. Huan Guo plotted figures for the Supplementary Information. Xianming Liu, Fangfei Zhang, and Tiannan Guo analyzed data with Spectronaut software. Jianwei Shuai, Yiming Li, Mingxuan Gao, Rongshan Yu, Donghui Guo, and Fangfu Ye discussed the algorithms. Jianwei Shuai and Jiahuai Han wrote the manuscript and supervised the project.

## Methods

### Training data for deep learning

*Escherichia coli* DH5a strain cells were washed three times with H2O and collected by centrifugation. Protein pellet was dissolved in 1%SDC/10 mM TCEP/40 mM CAA/Tris-HCl pH 8.5. Subsequently, 1% SDC was diluted with water to 0.5 %. The protein centration was measured with Pierce 660 nm protein assay reagent (Thermo). The trypsin (Sigma) was added with the ratio of 1:100 (trypsin: protein). The tubes were kept at 37 °C for 12-16 hours. The peptides were desalted with SDB-RPS StageTips. Peptides were dissolved in 0.1%FA (Formic Acid, 06440, Sigma) and analyzed by TripleTOF 5600 mass spectrometry (AB Sciex).

Peptides first bound to a 5 mm × 300 μm trap column packed with Zorbax C18 5-μm 300-Å resin (5065-9913, Agilent) using 0.1% (V/V) FA/2% ACN in H2O at 10 μl/min for 5 min, and then separated using 30, 45, 60, 120, 150, 180 or 240 -min gradient from 2-35% buffer B (buffer A 0.1% (V/V) FA, 5% DMSO in H2O, buffer B 0.1% (V/V) FA, 5% DMSO in CAN) on a 30 cm × 75 μm in-house pulled emitter-integrated column packed with Magic C18 AQ 3-μm 200-Å resin. The column temperature was kept at 50 °C by a column heater (PST_CHC-RC, Phoenix S&T) and a controller (PST-BPH-20, Phoenix S&T). For SWATH-MS, and MS1 scan records a 350-1250 m/z range for 250 ms and a 100-1800 m/z range was recorded for 33.3 ms in the high-sensitivity mode MS2 scan. One MS1 scan was followed by 100 MS2 scans, which covered a precursor m/z range from 400-1200. The variable windows of SWATH-MS were “399.5-409.9,408.9-418.9,417.9-427.4,426.4-436,435-443.6,442.6-450.8,449.8-458,457-464.8,463.8-471.1,470.1-476.9,475.9-482.8,481.8-488.6,487.6-494,493-499,498-504.4,503.4-509.3,508.3-514.3,513.3-519.2,518.2-524.2,523.2-529.1,528.1-534.1,533.1-539,538-543.5,542.5-548.5,547.5-553,552-558,557-562.5,561.5-567,566-571.5,570.5-576,575-580.5,579.5-585,584-589.5,588.5-594,593-598,597-602.5,601.5-607,606-611.1,610.1-615.6,614.6-620.1,619.1-624.6,623.6-628.6,627.6-633.1,632.1-637.6,636.6-642.1,641.1-646.6,645.6-651.1,650.1-655.6,654.6-660.1,659.1-665.1,664.1-669.6,668.6-674.5,673.5-679,678-684,683-688.5,687.5-693.4,692.4-698.4,697.4-703.3,702.3-708.7,707.7-713.7,712.7-719.1,718.1-724.5,723.5-729.9,728.9-735.3,734.3-740.7,739.7-746.5,745.5-751.9,750.9-757.8,756.8-763.6,762.6-769.5,768.5-775.3,774.3-781.2,780.2-787,786-793.3,792.3-800.1,799.1-806.4,805.4-813.1,812.1-820.3,819.3-827.5,826.5-835.2,834.2-843.3,842.3-851.4,850.4-859.9,858.9-868.9,867.9-878.4,877.4-888.3,887.3-899.1,898.1-910.3,909.3-922.9,921.9-936,935-949.5,948.5-963.4,962.4-978.7,977.7-994.9,993.9-1015.6,1014.6-1042.2,1041.2-1070.1,1069.1-1100.7,1099.7-1140.7,1139.7-1196.5”.

### Sample preparation and mass spectrometric analysis of L929 mouse datasets

Murine fibroblast L929 cells were harvested by scraping and centrifugation at 4 °C. L929 cells were lysed with 1% SDC/100 mM Tris-HCl pH 8.5, followed by sonication. The protein concentration was assayed using BCA. 10 μg proteins were reduced and alkylated using 10 mM TCEP/40 mM CAA. 1% SDC were diluted to 0.5% SDC using HPLC H2O, and trypsin was added at the protein: trypsin ratio of 50:1. Digestion was performed at 37 °C for 12 and 16 hours. The tryptic peptides were cleaned up using SDB-RPS StageTips before MS analysis.

Peptides were dissolved in 0.1% formic acid and analyzed by mass spectrometry in DDA and SWATH modes. MS analysis was performed on a TripleTOF 5600 (Sciex) mass spectrometry coupled to NanoLC Ultra 2D Plus (Eksigent) HPLC system. Peptides first bound to a 5 mm × 500 μm trap column packed with Zorbax C18 5-μm 200-Å resin using 0.1% (V/V) formic acid/2% acetonitrile in H2O at 10 μl/min for 5 min, and then separated from 2 to 35% buffer B (buffer A: 0.1% (V/V) formic acid, 5% DMSO in H2O, buffer B: 0.1% (V/V) formic acid, 5% DMSO in acetonitrile) on a 15 cm × 75 μm in-house pulled emitter-integrated column packed with Magic C18 AQ 3-μm 200-Å resin. For DDA, the 250-ms MS1 scan was performed in the range of 350-1250 m/z, and up to 20 most intense precursors with charge state 2-5 were isolated for fragmentation. MS/MS spectra were collected in the range of 100-1800 m/z for 100 ms. For SWATH-MS, a 100-ms survey scan (TOF-MS) which was collected in 350-1500 m/z was performed followed by 100 MS2 experiments with scan time 33 ms, which were collected in 100-1800 m/z. The 100 variable isolation windows of L929 dataset were the same as those of *E. coli* dataset acquired from TripleTOF 5600.

### The complete analysis workflow of Dear-DIA^XMBD^

The complete analysis workflow of Dear-DIA^XMBD^ mainly consists of identification and quantification processes. The workflow begins with a profile mzXML file and ends with a report file contained peptides and proteins. In the identification process, the raw files of mass spectrometry data were converted into profile mzXML files using MSConvert (V.3.0.19311), which were subjected to Dear-DIA^XMBD^ for generating pseudo-DDA mgf files (**Supplementary Fig. 3**). The mgf files were converted to mzXML files, which were analyzed with TPP (Trans-Proteomic Pipeline, Version 6.0.0) software. The mzXML files were subjected to database search using Comet (Version 2021.01) and X!Tandem (Version 2013.06.15.1, native and k-score) against an UniprotKB/Swiss-Prot database (**Supplementary Fig. 3**).

The pep.xml search results were validated and scored using PeptideProphet^51^ with parameters -p0.05 -l7 -PPM -OAdPE -dDECOY and combined by iProphet^52^ with parameters DECOY=DECOY. The Mayu.pl script (version 1.07) was used to determine iProphet probability corresponding to 1% peptide FDR. The peptide ions passing the 1% FDR were input into SpectraST^53^ for library building with CID-QTOF setting. The retention time of peptides in sptxt file was replaced with iRT time using spectrast2spectrast_irt.py script (downloaded from www.openswath.org), and iRT peptides used for retention time normalization were endogenous peptides. The sptxt file was made consensus nonabundant spectral library with the iRT retention time using SpectraST (**Supplementary Fig. 3**).

For the quantification process, the consensus sptxt files were converted to tsv using spectrast2tsv.py script which was then converted to TraML file with TargetedFileConverter tool integrated into OpenMS software (Version 2.6.0). OpenSWATH was run with options “-tr lib.os.pqp -threads 128 -sort_swath_maps - readOptions cacheWorkingInMemory -rt_extraction_window 600 - Scoring:TransitionGroupPicker:background_subtraction original -Scoring:stop_report_after_feature -1 -batchSize 0 -tr_irt irt.TraML - swath_windows_file win.os.tsv -out_osw”. An extended version of Pyprophet^54, 55^ (v2.1.10, Python 3.7, https://github.com/PyProphet) was employed for FDR estimation. 1% protein FDR at global level is applied in all analysis. The filtered results were input into TRIC^56^ software for cross-run alignment (**Supplementary Fig. 3**). The parameters in TRIC were set as followed: --method LocalMST --realign_method lowess_cython --max_rt_diff 60 --mst:useRTCorrection True --mst:Stdev_multiplier 3.0 --target_fdr 0.01 --max_fdr_quality 0.01 --verbosity 1 --alignment_score 0.0001 --in *.mzXML.tsv. The results were filtered at 1% global protein FDR. The final output file aligned.tsv contains the quantified peptide and protein information.

The TRIC results were used for protein inference and quantification. The proteins with proteotypic peptides were considered as “uniquely identified”. The peptides mapped to more than one protein entry were handled as followed. (1) The peptides shared with the proteins with proteotypic peptides were excluded from protein inference and quantification. (2) The peptides without evidence of unique protein mapping were considered as “from one protein representing the gene locus and expressed as the alphabetically first entry of the protein database (gene locus identification)”.^57^

### Parameters of software tools

The parameters of Comet and X!Tandem search engines are shown in **Supplementary Table 2**. The parameters of Dear-DIA^XMBD^, Spectronaut 14 (v14.10.201222.47784 and v14.9.201124.47784) and DIA-Umpire (v2.1.3) are shown in **Supplementary Table 3**, **Table 4**, and **Table 5**, respectively.

### Format conversion of benchmarked datasets

The .wiff raw data files were converted into profile and centroid mzXML format by msconvert.exe and qtofpeakpicker.exe from ProteoWizard (version 3.0.20039) package. The .raw files from Thermo Fisher Scientific mass spectrometer were converted into mzXML by msconvert.exe (ProteoWizard version 3.0.20039).

### Data preprocessing

Usually, DIA data contains a large number of background ion signals, which greatly increases the data redundancy and complexity. Thus, we applied several preprocessing algorithms to reduce the calculation consumption. In an MS1 isolation window, Dear-DIA^XMBD^ employs a fixed-width slider in MS1 retention time dimension to capture the local characteristics of DIA data. A slider contains a series of precursor ion spectra and the corresponding fragment ion spectra. Alignment of fragment XICs can be naturally resolved by using sliders in a single run. The fixed width of slider was set as 20, which is the length of XIC. Under the parallel mode, we moved the slider of all MS1 windows with stride of one and update the internal MS1 and MS2 spectra. By setting the appropriate width of slider, we assume that the peptides in a slider are recorded only once, and the chromatographic peaks of fragments from the same peptide show similar shapes.

Considering that precision of MS data in profile mzXML file as high as 10^−3^, Dear-DIA^XMBD^ applied the binning algorithm to truncate the precision MS data to low-precision values for spectrum analysis. In detail, we split 1.0 m/z to 30 bins with a truncated resolution of 0.03 m/z. The number of bins is a configurable parameter for the users. As a result, each spectrum in slider is represented by a fixed-length vector after data binning. For instance, if the maximum value of m/z is specifically defined as 1200, the spectrum will be represented by a vector of length 36,000. An index of the vector corresponds to m/z of an ion, and the vector value at that index is equal to the ion intensity. If an ion is not recorded in a spectrum, its intensity will be replaced by zero. The binning algorithm starts from the zero value of initial vector. Then the ion intensity is accumulated to the vector values of the corresponding index. Both MS1 and MS2 spectra are handled by this binning method.

Because the signals of precursors and fragments are submerged in a large number of background ion signals, the binned DIA data is still complicated. It is important to remove the background ions with an extremely low signal-to-noise ratio (SNR). Furthermore, in SWTAH workflow, MS1 and MS2 scan times are configured to 250 milliseconds and 33 milliseconds, respectively. The difference in scan time causes the SNR of MS1 spectrum to be higher than that of MS2 spectrum. Therefore, Dear-DIA^XMBD^ adopts different methods to filter the background ions in MS1 and MS2 spectra.

For each MS1 spectrum, a peak-finding algorithm is applied to detect peaks with high SNR in m/z dimension. Those detected peaks are probably from the true signals of peptides, rather than background ions. For the detected peaks, a deisotoping algorithm is then used to find the isotopic clusters and to calculate the ion charge of the first isotopic peak. The ions that are able to determine the ion charges are regarded as the candidate precursors in a slider. Dear-DIA^XMBD^ stores XICs, charges and the binning m/z indexes of the candidate precursor ions.

Furthermore, the background ions in MS2 spectra are handled by setting two filter conditions of SNR of XICs. The first requirement is that the number of non-zero values of XIC must be greater than five. The second condition requires that the ratio of the maximum value to the non-zero minimum value of XIC should be larger than four. All fragments satisfying these conditions are treated as the candidate fragments in a slider, and their m/z and XICs are stored for the next processing.

### Feature extraction of fragment XICs

We feed fragment XICs into the encoder network and store the output of encoder as the representation of XICs. The deep neural network is written by Python3.6 on MXNet deep learning framework and trained on NVIDIA GeForce GTX 1080Ti GPU.

### Architecture and training process of variational autoencoder

Autoencoders are important unsupervised learning models for data dimensionality reduction and feature extraction. Their learning objectives perform the following mapping:

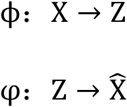

where Z represents the features of input data. The encoder network ϕ maps input data X to Z, and then the decoder network φ reconstructs Z to 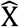. The input data X is the XICs of fragments in a slider. The learning objective of autoencoder is to make 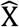 as close to X as possible.

The common encoder and decoder are designed as a stack of fully connected neural networks, which are simple with high computing speed. In order to achieve better performance on feature extraction task, we referenced the four-branch networks idea of GoogLeNet^58^ structure and constructed a four-branch of fully connected VAE neural networks. In the networks, we set the number of neurons of fully connected layers to be equal to the channel size of inception block.

The network structure of encoder and decoder presents mirror symmetry (**Supplementary Fig. 1**). The encoder network is a four-branch network, and each branch consists of fully connected (FC) layers. The first branch network contains a fully connected 384-dimensional layer, followed by a dropout layer. The second branch network includes two fully connected layers with the dimensions of 192 and 384, and a dropout layer. The third branch network includes two fully connected layers with the dimensions of 48 and 128, and a dropout layer. The fourth branch only contains a fully connected 128-dimensional layer (**Supplementary Fig. 1**).

The 20-dimensional input vector of encoder network is fragment XIC. The output vectors of the four-branch networks are catenated by the appending operation at the end. The encoder network outputs two 16-dimensional vectors, one for the standard deviation (*σ^2^*) and the other for the mean value (*μ*). The mean vector represents the latent features of the input data. Then, 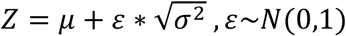 is fed to the decoder network, where *ε* is a random value sampled from Gaussian distribution (**Supplementary Fig. 1**).

Then the 16-dimensional vector *Z* is fed to the decoder network consisting of four-branch network. For the decoder network, the first branch network contains a fully connected 384-dimensional layer, followed by a dropout layer. The second branch network includes two fully connected layers with the dimensions of 384 and 192, and a dropout layer. The third branch network includes two fully connected layers with the dimensions of 128 and 48, and a dropout layer. The fourth branch only contains a fully connected 128-dimensional layer. The output vectors of the four-branch networks are catenated by the appending operation at the end, and the size of catenation vector is 384+192+48+128=752. Then, the 752-dimensional catenation vector is fed to a 20-dimensional fully connected layer. Finally, the decoder network outputs a 20-dimensional vector as the reconstructed data of the input vector (**Supplementary Fig. 1**).

To train the variational autoencoder, we input the anchor, positive and negative XICs (*X_a_*, *X_p_*, *X_n_*) to the encoder network, respectively (**Fig. 2a**). The encoder network outputs the latent features (*μ_a_*, *μ_p_*, *μ_n_*) and the variance values 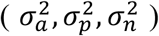 corresponding to the anchor, positive and negative XICs, respectively. Then, *Z_a_*, *Z_p_* and 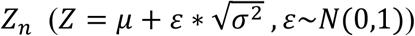 are fed to the decoder network, respectively. The decoder network reconstructs *Z_a_*, *Z_p_* and *Z_n_* to 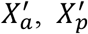 and 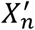, respectively. Here we calculate the objective functions of classical VAE of anchor, positive and negative XICs, respectively. The objective function of the classical VAE is defined by the following equations:

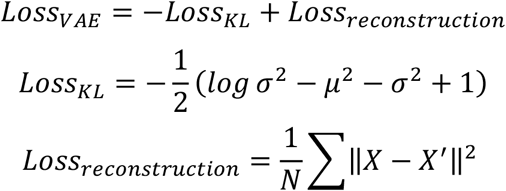

The *Loss_VAE_* of anchor, positive and negative XICs is defined to 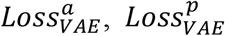 and 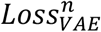. Then, we calculate the triplet loss by using *μ_a_*, *μ_p_* and *μ_n_*. The triplet loss is defined by the following equation:

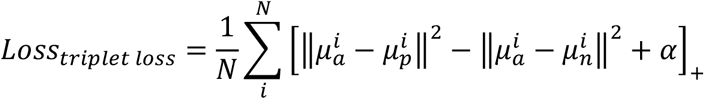

where *α* is a margin parameter which is set to 1. In addition, *‖*‖^2^* presents the square of Euclidean distance. Finally, we combine the VAE loss and the triplet loss as the final optimized function *Loss_total_*:

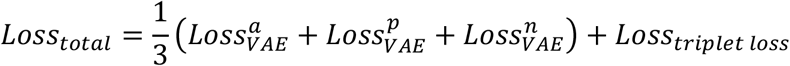

In the above training process, the anchor, positive and negative XICs are input to the encoder network, respectively. The encoder network outputs the latent features of each input fragment XIC, whether it is anchor, positive or negative. Therefore, when making prediction, the input data of the trained neural network model is all fragment XICs in a slider, rather than a single labeled XIC.

We employed 6 common deep learning optimizers (Adadelta, Adagrad, Adamax, Nadam, SGD and Adam) to optimize our model. By comparing the loss function curves (**Supplementary Note 1 Fig. 1**) of different optimizers, we decided to use Adam (adaptive moment estimation) to optimize our model to find more peptides. The update rules of Adam optimizer are defined by the following formula:

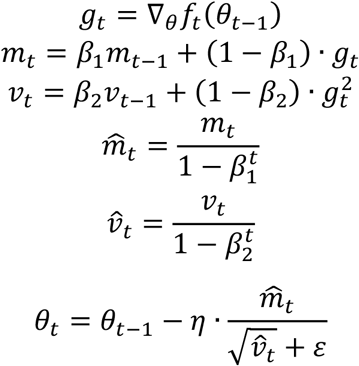

where *f_t_*(*θ*_*t*-1_) is the loss function, *g_t_* is the gradient of the parameter *θ*, and *η* is the learning rate. The default value of *η* is 0.001. *β_1_* and *β_2_* are the parameters in the algorithm, generally *β_1_ = 0.9* and *β_2_ = 0.999*. *m_t_* and *v_t_* are the first-order and the second-order moment estimation of the gradient, respectively. 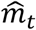 and 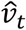 are corrections to *m_t_* and *v_t_*, respectively, which can be approximated as an unbiased estimate of the expectation. When *ε* = 10^−8^, the zero denominator can be avoided.

### Triplet dataset generation

When the triplet loss is introduced into the model, we need to generate the triplet data to train neural network. The training data comes from the results quantified by OpenSWATH. We stored XICs of the first six high-intense fragments of quantified peptides. Then, we randomly chose two fragment XICs from the same peptide as anchor and positive data. The negative data were randomly selected from the different peptides. Finally, we combined the anchor, positive and negative data into triple data.

### Architecture and training process of CNN classifier

We applied CNN classifier to calculate the similarity of hit fragments which matched in-silico peptide. Since the first 6 fragment ions were usually selected to observe their similarity during manual check, we fed the XICs of first 6 ions in hit fragments into CNN classifier. We used 1 and 0 to label fragment ions belonging to the same peptide and different peptide, respectively.

The length of each fragment XIC is 20, so that the input matrix of CNN contains 6 rows and 20 columns (**Supplementary Fig. 2**). The CNN classifier consists of four-branch (1-2-2-1) convolutional layers, which is the same with Inception block of GoogLeNet. The output feature maps of the four-branch networks are catenated in channel dimension. The result of catenation is flattened into a vector, which is fed to a 512-dimensional fully connected (FC) layers. The last layer reports the similarity score, which locates between 0 to 1 (**Supplementary Fig. 2**). The loss function of CNN classifier is the Binary Cross Entropy (BCE) function, which is defined by the following equation:

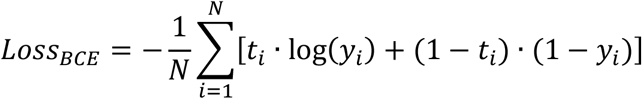

where *t_i_* and *y_i_* represent the label of input data and the output of CNN classifier, respectively. *N* is the number of input matrixes. We used Adam optimizer with default parameters to train CNN classifier and treated the outputs of CNN classifier as the similarity scores of input fragment groups.

### PIndex querying algorithm

In order to match fragments with precursors, we developed PIndex querying algorithm based on the inverted index algorithm. PIndex starts with *in silico* digestion of protein FASTA database and then generates the *in silico* digested peptide information sets which contain the charge of precursor, the m/z of precursor, and the m/z list of fragments. We allocated the unique index to each information set. Obviously, we can query the *in silico* precursors and fragments by using the peptide indexes.

Next, we created the inverted index table between the peptide indexes and the *in silico* digested peptides. The inverted querying process includes two parts, one is to map precursors to peptide indexes and the other is map fragments to peptide indexes. Precursor query maps the precursor identifiers, including m/z and charge, to peptide index set which is named Index1. Fragment query maps the fragment m/z to peptide index set which is named Index2. We calculated the intersection of Index1 and Index2, then we can obtain the peptides which were hit by both fragments and precursors. Querying the same peptide index indicates that the precursor and fragments come from the same peptide.

### Two-stage clustering method

In the first k-Means clustering, we obtained numerous fragment ion combinations. These combinations were matched with precursors by using PIndex querying. However, affected by the interference signal, there are still some fragment ions in the clustering results that did not match the precursors. Therefore, we removed the precursor-fragment pairs in the first clustering, and used k-Means to cluster the remaining fragment ions again to improve the usage of fragment ions.

### MS1 recalibration

When the MS is not calibrated for long, the masses will often exhibit systematic shifts. The proper calibration can improve identification, alignment and quantification. We referenced mzRecal^40^, a universal MS1 recalibration method using high confidence peptides as internal calibrants, to improve the performance of Dear-DIA^XMBD^. We selected the peptides with X!Tandem expected values less than 0.001 as potential calibrants, and then use the following mzRecal formula to calibrate the MS1 m/z:

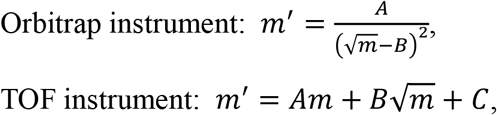

where *m*′ is the calibrated m/z, m is the experimental m/z. Parameters A, B and C can be calculated by curve fitting method.

